# A novel host-encoded protein, ETP2, plays an important role in endosymbiont division in the trypanosomatid *Angomonas deanei*

**DOI:** 10.1101/2025.04.22.649992

**Authors:** Anay K. Maurya, Lawrence Rudy Cadena, Georg Ehret, Eva C. M. Nowack

## Abstract

A single β-proteobacterial endosymbiont, *Candidatus* Kinetoplastibacterium crithidii, resides in the cytosol of the trypanosomatid *Angomonas deanei*, and divides at a defined stage of its host’s cell cycle. This endosymbiont has a highly reduced genome of 0.8 Mb and, notably, has lost most essential genes involved in bacterial cell division, resulting in a loss of division autonomy. It has been previously demonstrated that a host-encoded dynamin-like protein, ETP9, plays an indispensable role in the division of the endosymbiont. In this study, we identified a second nucleus-encoded component of the endosymbiont division machinery, termed ETP2, currently annotated as a ‘hypothetical protein’. We observed that ETP2 localizes in a cell cycle-dependent manner at the bacterial division site, alongside the bacterium-encoded FtsZ. Furthermore, we demonstrated that ETP2 plays an important role for timely endosymbiont division and segregation, as a large fraction of cells in an *etp2* deletion mutant cell line exhibited either long, filamentous endosymbionts accompanied by severely distorted host cells or host daughter cells lacking endosymbionts. *In silico* analyses revealed that ETP2 is found exclusively in endosymbiont-harboring trypanosomatids and is most likely an intrinsically disordered protein. Collectively, our data suggests that ETP2, alongside the previously described ETP9, is an integral component of the endosymbiont division machinery. This finding highlights the evolution of a complex host-derived molecular mechanism that exerts tight control over its endosymbiont without requiring gene transfers from the bacterium.

## Introduction

The transformation of a free-living bacterium into a fully integrated organelle involves extensive structural, physiological, and genetic changes (Bhattacharya et al., 2007; Roger et al., 2017). Although mitochondria and plastids originated from bacteria, over a billion years of co-evolution with their host have led to a situation where the organelles and the surrounding cell can no longer be viewed as independent entities. Instead, the organelles’ proteome composition, metabolic activity, division timing, positioning within the cell, and other factors are predominantly regulated by the genetic instructions of the nucleus. More recently established endosymbiotic relationships offer a unique opportunity to observe intermediate stages in the process by which a prokaryote becomes genetically integrated into a eukaryotic cell (Sørensen et al., 2024). As a result, studying the molecular mechanisms that govern these host-endosymbiont interactions holds significant potential for uncovering the scenarios and molecular processes behind organelle formation.

Interestingly, some protists that have developed vertical endosymbiont transmission from one host generation to the next, exhibit strict control over the number of endosymbionts per host cell. Examples include the cercozoan *Paulinella chromatophora*, which houses two cyanobacterium-derived photosynthetic “chromatophores” (Nowack, 2014), the haptophyte *Braarudosphaera bigelowii*, which contains a single cyanobacterium-derived nitrogen-fixing “nitroplast” (Coale et al., 2024), and the trypanosomatid *Angomonas deanei*, which contains a single β-proteobacterial endosymbiont that supplies its host with diverse metabolites and co-factors (Mundim and Roitman, 1977; Alves et al., 2011; Alves et al., 2013; Harmer et al., 2018). Intriguingly, this endosymbiont, *Ca.* Kinetoplastibacterium crithidii, consistently divides at a specific time point in the host cell cycle (Motta et al., 2010). Presence of *Ca.* K. crithidii seems to be obligate in some *A. deanei* strains including *A. deanei* ATCC-PRA 265, the strain investigated in this study (Maurya et al., 2025). However, in other *A. deanei* strains, artificially aposymbiotic cells that lost the endosymbiont can be generated through antibiotic treatment when metabolites normally provided by the endosymbiont are externally provided (Mundim and Roitman, 1977).

The *Ca.* Kinetoplastibacterium endosymbiont was apparently acquired approximately 40-120 million years ago (MYA) by a common ancestor of the trypanosomatid subfamily Strigomonadinae, to which *A. deanei* belongs (Du et al., 1994). Thus, it is also present in species of the other genera in this subfamily, *Strigomonas* and *Kentomonas* (Teixeira et al., 2011; Votýpka et al., 2014). All *Ca.* Kinetoplastibacterium endosymbionts have highly reduced genomes (∼0.8 Mbp) (Alves et al., 2013; Silva et al., 2018). Notably, this reductive genome evolution included the loss of most genes encoding essential components of the bacterial division machinery (Motta et al., 2013; Maurya et al., 2025). However, the gene encoding the central cell division protein FtsZ has been retained. FtsZ is a polymer-forming GTPase that forms in free-living bacteria as well as plastids and some mitochondria the so-called Z-ring at mid cell at the inner face of the inner membrane (Margolin, 2005; Leger et al., 2015). In bacteria, the Z-ring serves as a scaffold for the assembly of the machinery that synthesizes the division septum. In plastids, the Z-ring forms a complex with proteins in the inner and outer envelope membrane that confers topological information to the outer face of the outer envelope membrane and recruits a soluble dynamin-like protein (DLP) that forms a contractile ring structure around the plastid. Reciprocal interactions between the Z-ring inside and the DLP ring outside result in a concerted constriction of the division machinery responsible for plastid fission (Wang et al., 2017).

Trypanosomatids that thrive in their natural habitat as parasites, mostly in the digestive tract of insects, are capable of growing axenically in culture to high densities, can be cryopreserved, and are particularly well-suited for genetic manipulation (Clayton, 1999; Lukeš et al., 2018; Yagoubat et al., 2020; Kostygov et al., 2021). Hence, the Strigomonadinae represent an optimal model to investigate the molecular mechanisms establishing nuclear control over endosymbiont division. The development of genetic tools for *A. deanei* together with the mass spectrometric characterization of the endosymbiont proteome recently enabled the identification of seven <u>e</u>ndosymbiont-targeted host <u>p</u>roteins (ETPs) (Morales et al., 2016; Morales et al., 2023). One of these proteins is the DLP ETP9 that apparently forms a contractile ring around the endosymbiont-division site (ESDS) -likely on the outer face of the endosymbiont’s outer membrane-and takes part in endosymbiont division in a manner analogous to the division of mitochondria and plastids (Morales et al., 2023; Maurya et al., 2025). *A. deanei* is a diploid organism, haploid stages are not known. Attempts to generate homozygous ETP9 knockouts in which both *etp9* alleles are replaced by a resistance marker gene, proved unsuccessful, suggesting that this nucleus-encoded protein is essential for the organism’s survival. ETP9 knockdown (KD) experiments demonstrated that despite the formation of Z-ring structures in ETP9-depleted cells, endosymbionts are division-impaired, forming long, filamentous structures, within significantly distorted host cells (Maurya et al., 2025). How ETP9 is recruited to the ESDS, if it acts alone or is part of a more complex division machinery, and if nucleus-encoded endosymbiont division factors are physically linked to the bacterial Z-ring is currently unknown.

Notably, ETP9 is not the sole ETP that was found to localize at the ESDS, as ETP2 (annotated as a ‘hypothetical protein’) and ETP7 (containing a predicted phage-tail lysozyme domain) exhibited similar localizations (Morales et al., 2023). Here, we investigated the role of ETP2 in *A. deanei*. By using a combination of microscopy and knockout/KD approaches, we observed that ETP2 shows a cell cycle-dependent localization at the ESDS and fulfills an important role in endosymbiont division and segregation into host daughter cells. *In silico* analyses found ETP2 restricted to the Strigomonadinae and is likely an intrinsically disordered protein. Together our data suggests that ETP2 evolved as a component of the endosymbiont division machinery in *A. deanei*, exhibiting a tight control of the host over its bacterial endosymbiont.

## Results

### Recombinant eGFP-ETP2 is functional and consistently localizes at the ESDS

Previously, we observed that ETP2 (GenBank accession CAD2221027.1) fused at its N-terminus to the green fluorescent protein eGFP, overexpressed from the *δ-amastin* locus (Δ*δ-ama^egfp-etp2^*) localizes at the ESDS (**Fig. 1A** and (Morales et al., 2023)). To verify that recombinant ETP2 expressed from its endogenous locus showed the same subcellular localization, we tagged ETP2 endogenously (Δ*etp2^egfp-etp2^/etp2)* in a parental strain expressing the endosymbiont marker mScarlet-ETP1 (Δ*γ-ama^mS-etp1^*; see (Morales et al., 2023)) or in the wildtype (Wt) background. Epifluorescence microscopy revealed that in both cell lines, the endogenously tagged ETP2 showed the same localization as when overexpressed, although the eGFP fluorescence signal was weaker (**Fig. 1B**, micrographs 1 and 2). Cells in which both alleles of *etp2* were replaced with the recombinant version (Δ*etp2^egfp-etp2^/*Δ*etp2^egfp-etp2^*) showed an enhanced eGFP fluorescence signal at the ESDS (**Fig. 1C**; for confirmation of the cell line by PCR and Southern blot see **Fig. S1**). Furthermore, no unusual phenotype was observed in these cells, suggesting that its N-terminal fusion to eGFP neither affects ETP2 function nor subcellular localization.

**Figure 1:**
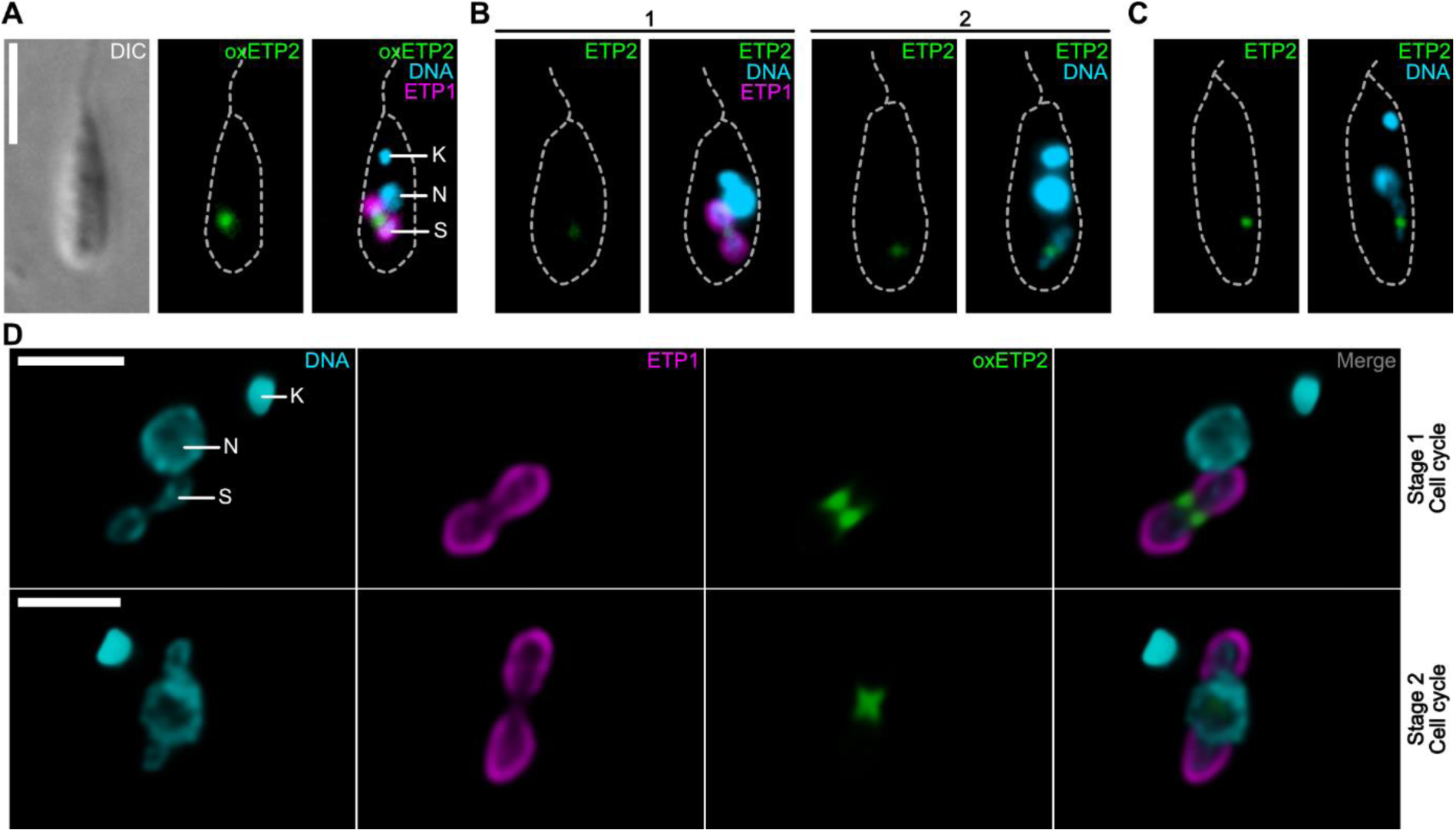
Subcellular localization of recombinant ETP2 in *A. deanei*. **A-C:** Epifluorescence microscopic analysis of subcellular localization of (A) eGFP-ETP2 overexpressed (oxETP2) from the *δ-amastin* locus in a cell co-expressing the endosymbiont marker mScarlet-ETP1; (B) eGFP-ETP2 expressed from its endogenous locus in the background of a cell expressing mScarlet-ETP1 (micrograph 1) or the Wt background (micrograph 2); and (C) eGFP-ETP2 expressed from both endogenous loci. For each image set, the eGFP channel alone (left) and the overlay of channels (right) for Hoechst 33342-stained DNA, eGFP, and (where relevant) mScarlet fluorescence are shown (A-C). As illustrated in panel A, the broken grey lines show cell outlines as seen by differential interference contrast (DIC) light microscopy. Scale bar is 5 µm. **D:** Deconvoluted fluorescence signals from confocal microscopy of eGFP-ETP2 overexpressing cells (same cell line as in A). Individual channels as well as the overlay of signals from channels for Hoechst 33342, mScarlet, and eGFP are shown. Upper row, early stage in endosymbiont division; lower row, later stage. Scale bar is 2 µm. Abbreviations: K, kinetoplast (= network of concatenated mitochondrial DNA); N, nucleus; S, endosymbiont.

In optical transects of the ESDS by confocal microscopy, (overexpressed) eGFP-ETP2 forms two patches at both sides of the endosymbiont envelope (**Fig. 1D**, cell cycle stage 1). These patches appear to be part of a ring-shaped structure surrounding the ESDS as observed when moving through the Z-stacks (**Movie S1**). As the bacterium elongates and constricts in the middle, during bacterial division, the two eGFP-ETP2 patches seen in the optical transects meet in the middle of the ESDS, slightly cupping the newly forming bacterial cell poles, resulting in an x-like structure (**Fig. 1D**, cell cycle stage 2 and **Movie S2**).

### Endosymbiont-encoded FtsZ and host-encoded ETP2 show a cell cycle-dependent localization at the ESDS

Previously, reconstruction of cell cycle stages of *A. deanei* had shown that the bacterium-encoded FtsZ and nucleus-encoded ETP9 exhibited a cell cycle-dependent localization at the ESDS (Maurya et al., 2025). To test whether ETP2 displayed a similar localization dynamic, we visualized FtsZ by immunofluorescence assay (IFA), and eGFP-ETP2 and the endosymbiont marker mScarlet-ETP1 by their autofluorescence. Since neither protocol for live-cell imaging nor synchronization of *A. deanei* cultures are available, we reconstructed division stages based on cell morphologies from images of >1000 fixed cells (**Fig. 2**).

**Figure 2:**
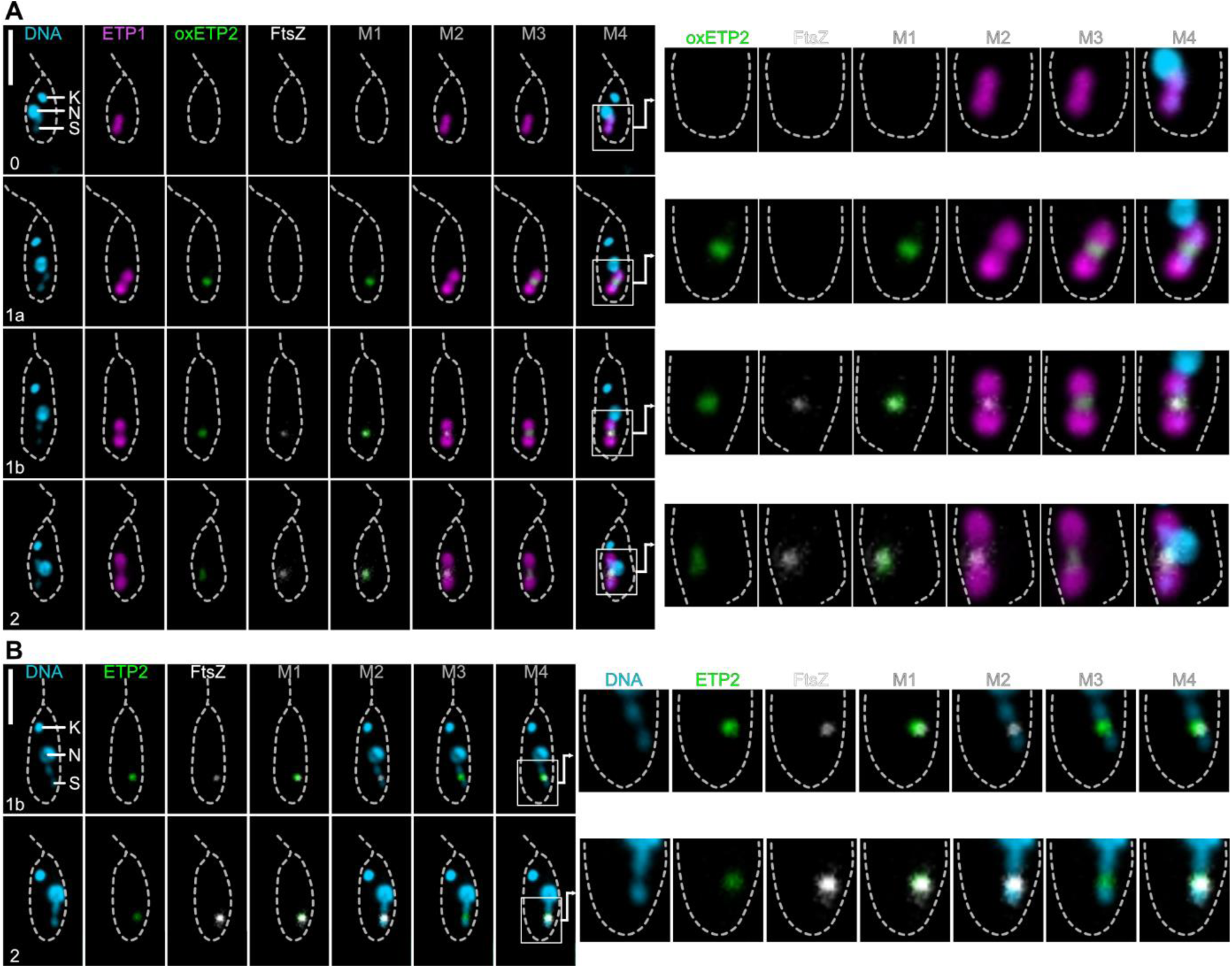
Subcellular localization of endosymbiont-encoded FtsZ and host-encoded recombinant ETP2 throughout early cell cycle stages. Subcellular localization of eGFP-ETP2 and FtsZ in mid-log phase cells co-expressing mScarlet-ETP1 (Δ*γ-ama^mS-etp1^*) and eGFP-ETP2 [overexpressed (Δ*δ-ama^egfp-etp2^*) in **A**; and endogenously tagged (Δ*etp2^egfp-etp2^*/Δ*etp2^egfp-etp2^*) in **B**] throughout early cell cycle stages as analyzed by epifluorescence microscopy. eGFP-ETP2 (green) and mScarlet-ETP1 (magenta) were detected by their autofluorescence, FtsZ (white) by IFA, and DNA (cyan) by Hoechst 33342 staining. Scale bars are 5 µm. Merges in A: Merge1, overlay of eGFP and FtsZ signals; Merge2, mScarlet and FtsZ; Merge3, mScarlet and eGFP; Merge4, all four channels. Merges in B: Merge1, overlay of eGFP and FtsZ signals; Merge2, FtsZ and Hoechst33342; Merge3, eGFP and Hoechst33342; Merge4, all three channels. Abbreviations are the same as in Fig. 1. Numbers on images show cell cycle stages compared to **Maurya et al., 2025**. White boxes in the left panels indicate areas of detailed images in the right panels.

For the eGFP-ETP2 overexpressing cell line (Δ*δ-ama^egfp-etp2^*; **Fig. 2A**), initially neither FtsZ nor eGFP-ETP2 fluorescence signals were observed (stage 0). Then, eGFP-ETP2 appears at the ESDS (stage 1a) followed by FtsZ, where they colocalize (stage 1b). During stage 2, the endosymbiont elongates with both FtsZ and ETP2 remaining co-localized at the ESDS. At later stages, the signal disappears in most cells, and only few cells show diffuse or inconsistent fluorescence signals for FtsZ and/or eGFP-ETP2. For the endogenously tagged ETP2 (Δ*etp2^egfp-etp2^/*Δ*etp2^egfp-etp2^*; **Fig. 2B**), the localization patterns appear similar, however, it seems that both proteins, eGFP-ETP2 and FtsZ, reach the ESDS at a similar time point. Hence, the apparently early arrival of the overexpressed eGFP-ETP2 before FtsZ (stage 1a) might rather reflect its better detectability than actual early arrival and a clear order of arrival cannot be unambiguously established.

### Generation of a homozygous *etp2* deletion mutant is viable and results in severe division phenotypes

To study the cellular function of ETP2, we initially generated a heterozygous *etp2* deletion mutant cell line in which one allele of *etp2* was replaced by the neomycin phosphotransferase gene, *neo^R^*, (Δ*etp2^neo^*/*etp2*; **Fig. 1S**). This mutant showed no noticeable phenotype. Several attempts to replace the remaining allele of *etp2* by a hygromycin phosphotransferase gene, *hyg^R^*, failed. However, we obtained a homozygous deletion mutant cell line, in which the second allele was disrupted by insertion of a *hyg^R^*-containing cassette into the *etp2* open reading frame (ORF) resulting in a heavily truncated ETP2 protein (Δ*etp2^neo^*/Δ*etp2_246-491_^hyg^*; **Fig. S1**). Both, heterozygous and homozygous *etp2* deletion mutants were generated not only in the symbiotic but also an aposymbiotic *A. deanei* strain (strains ATCC PRA-265 and ATCC 30969, respectively). All cell lines generated were verified by PCRs and Southern blot analysis (**Fig. S1**). Interestingly, in the symbiotic homozygous *etp2* deletion mutant cell line, >50% of the cells displayed either long, filamentous endosymbionts accompanied by severely distorted host cells or loss of the endosymbiont (**Fig. 3A-C**). This distortion was not witnessed in the aposymbiotic homozygous *etp2* deletion mutant cell line where cells did not differ morphologically from Wt cells (**Fig. S2**), suggesting that the function of ETP2 is confined towards the endosymbiont, with the resulting distorted host morphology caused by downstream effects of impaired endosymbiont division. This distortion of host cells was accompanied by the occurrence of multiple kinetoplasts and flagella, and notably, a singular and enlarged nucleus in most cases, suggesting that both kinetoplast and nuclear DNA replication remain active yet only segregation of the kinetoplast is fulfilled. However, ∼48% of the cells appeared ‘normal’ retaining a singular, morphotypical endosymbiont (**Fig. 3B-C**). Upon closer examination, a number of morphologically ‘normal’ host cells appear in the final stages of cytokinesis, attached only by an apparently undivided endosymbiont localized at the posterior end of the cells (**Fig. 3B**, white arrowhead). Additionally, some cells lacking endosymbionts are noted in the final stages of cytokinesis (**Fig. 3B**, black arrowhead), and possibly completing it without fulfilling complete segregation of the endosymbiont, explaining the high population of aposymbiotic cells in the homozygous mutant. Hence, the disruption of the second *etp2* allele is not entirely lethal, suggesting that although now error prone, successful endosymbiont division is still possible. This is reflected also by the slower growth of the *etp2* deletion mutant compared to Wt cells (**Fig. 3D**).

**Figure 3:**
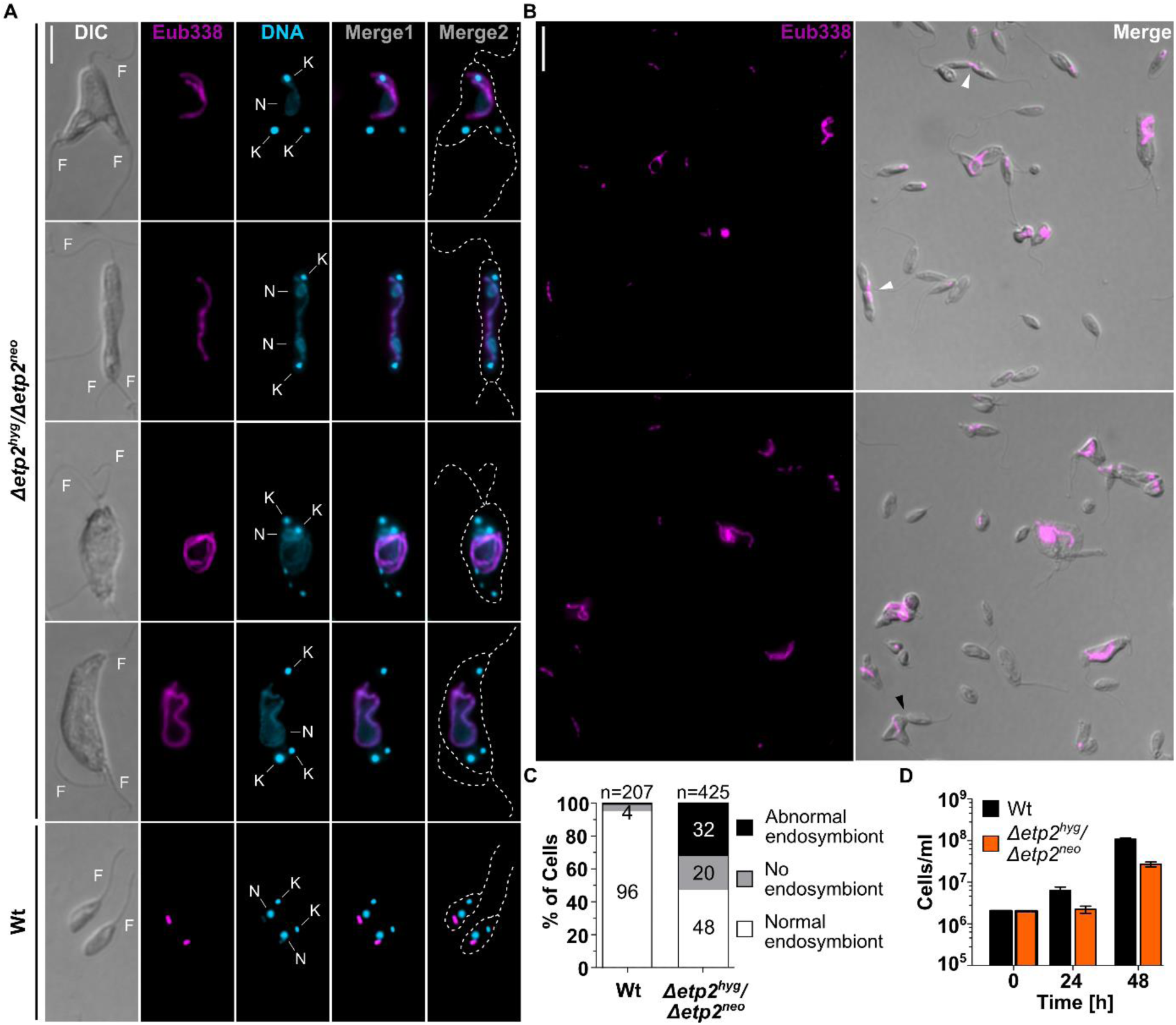
Characterization of homozygous *etp2* deletion mutants in symbiotic *A. deanei*. **A:** Fluorescence *in situ* hybridization (FISH) micrographs of homozygous *etp2* mutant cells. Wt *A. deanei* cells are shown as a control. Merge1, superposition of signals from Hoechst 33342-stained DNA (cyan) and the Cy3-Eub338 FISH probe directed against the bacterial 16S rRNA (magenta). Merge2, superposition of Hoechst 33342 and Cy3 signals on cell outlines as seen by DIC (broken grey lines). Abbreviations: K, kinetoplast; N, nucleus; F, flagellum. Scale bar is 5 µm. **B:** Overview micrographs of Cy3-Eub338 FISH-stained homozygous *etp2* mutant cells showing the occurrence of cells with abnormal, normal, or lacking endosymbionts. White arrowheads indicate host cells during the final stages of cytokinesis, however, attached by an undivided endosymbiont at the posterior end. Black arrowhead indicates a host cell in the final stage of cytokinesis lacking an endosymbiont. Scale bar is 10 µm. Note the multiple occurrences of cells lacking an endosymbiont. **C:** Quantification of division phenotypes in the *etp2* mutant compared to Wt cells. **D:** Cell counts of symbiotic Wt and *etp2* deletion mutant cells at 24 h and 48 h after inoculation (0 h). Plotted are mean and standard deviation from three biological and for each three technical replicates.

### KD of ETP2 results in elongated endosymbionts in severely distorted host cells

To confirm by an independent method that the observed division phenotype in homozygous *etp2* deletion mutants is caused by the lack of ETP2, we used a KD approach based on morpholino antisense oligos (MAOs).

For this, we transfected symbiotic and aposymbiotic *A. deanei* Wt cells with MAOs designed against *etp2* mRNA (MAO*_etp2_*) (**Fig. 4A**). Hybridization of MAOs with the 5’ UTR of a specific mRNA results in the repression of its translation (Summerton and Weller, 1997). Consistent with the phenotype observed in the *etp2* deletion mutants (**Fig. 3**), 24 h post-transfection, we observed a clear reduction in growth and the formation of long, filamentous endosymbionts in distorted host cells in the symbiotic strain (**Fig. 4B-C**), whereas in the aposymbiotic strain no noticeable changes in cell morphology were observed (**Fig. 4E**). However, a slight reduction in growth was observed in the aposymbiotic strain following transfection with MAO*_etp2_* compared to the water control, which might reflect off-target effects (**Fig. 4B**). For better visualization of the endosymbionts, cells expressing the endosymbiont marker mScarlet-ETP1 were also transfected with MAO*_etp2_* resulting the same filamentous endosymbionts (**Fig. 4D**; for overview images see **Fig. S3**).

**Figure 4:**
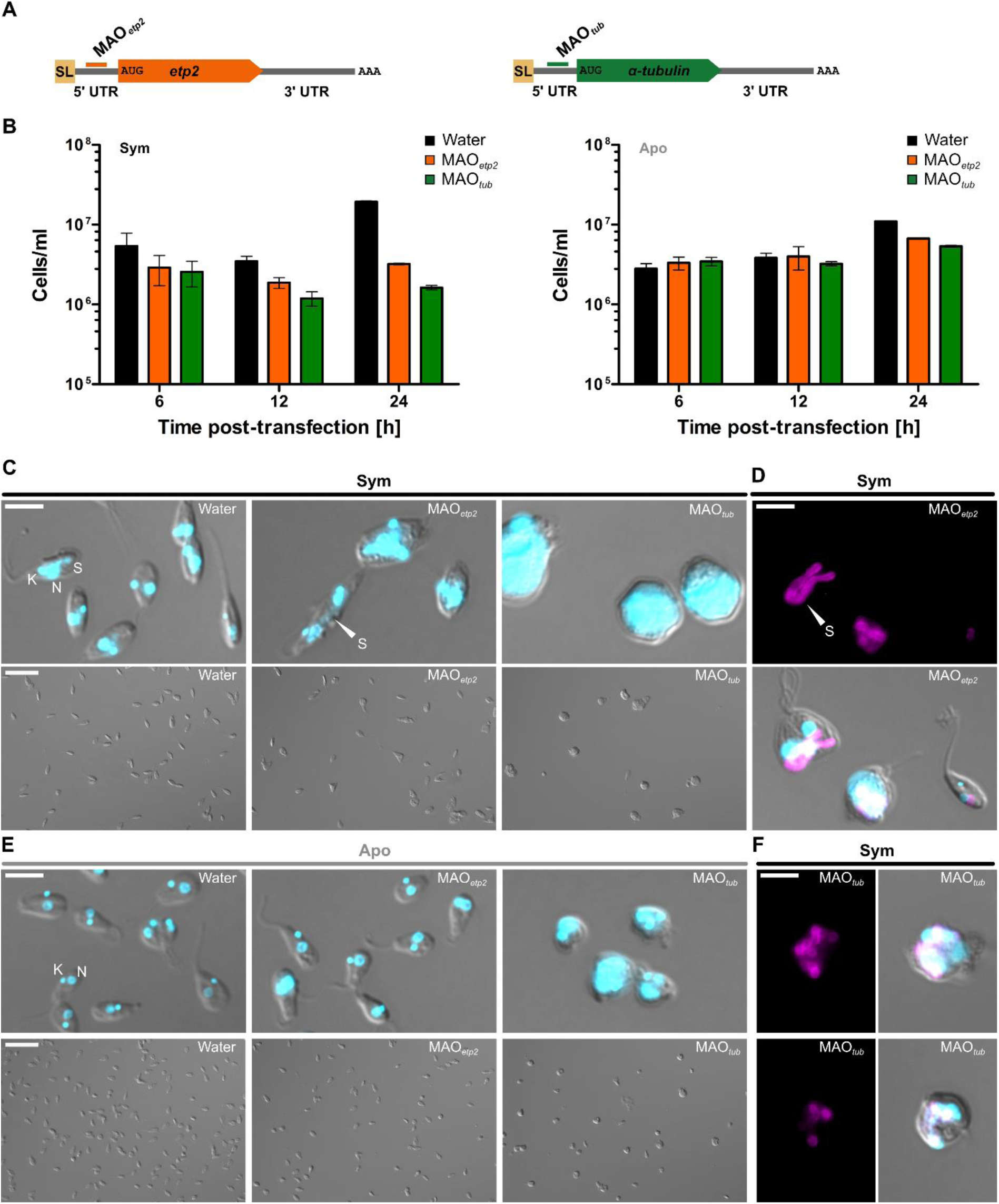
KD of ETP2 in symbiotic and aposymbiotic *A. deanei* strains. **A:** Schematic representation of MAO binding sites on the target mRNA of *etp2* and *α-tubulin*. **B:** Cell densities of cultures of the symbiotic and aposymbiotic *A. deanei* strains 6 h, 12 h, and 24 h post-transfection with water, MAO*_etp2_* or MAO*_tub_*. Plotted are mean and standard deviation from three technical replicates. **C:** Micrographs of the symbiotic cells 24 h post-transfection for each treatment. A detail micrograph (merge of DIC and Hoechst 33342, upper panel) and overview micrograph (DIC, lower panel) are shown. **D:** Micrographs of symbiotic cells expressing the endosymbiont marker mScarlet-ETP1 24 h post-transfection with MAO*_etp2_*. The mScarlet channel alone (upper panel) and merge of DIC, Hoechst 33342, and mScarlet channels (lower panel) are shown. (For overview images, **see Fig. S3**). **E:** Micrographs of aposymbiotic cells 24 h post-transfection for each treatment. A detail micrograph (merge of DIC and Hoechst 33342, upper panel) and overview micrograph (DIC, lower panel) are shown. **F:** Micrographs of symbiotic cells expressing the endosymbiont marker 24 h post-transfection with MAO*_tub_*. The mScarlet channel alone (left panel) and merge of DIC, Hoechst 33342, and mScarlet channels (right panel) are shown. Abbreviations: K, kinetoplast; N, nucleus; S, symbiont; SL: splice leader, and UTR: untranslated region. Arrowhead highlights the symbiont. Scale bars are 5 µm (for detail) and 25 µm (for overview micrographs).

Importantly, symbiotic as well as aposymbiotic cells mock treated with water showed the highest cell numbers and no aberrant cell morphologies 24 h post-transfection (**Fig. 4B-C, E**); while cells transfected with MAO*_tub_* against α-tubulin, as a positive control, exhibited the expected effects (**Fig. 4B-C; E**), with a markedly reduced growth and formation of roundish cell morphologies comparable to the “Fat Cells” phenotype described in *Trypanosoma brucei* following RNAi against α-tubulin (Ngô et al., 1998) and our previous results in *A. deanei* (Maurya et al., 2025). Furthermore, a massive accumulation of DNA-containing compartments was observed following α-tubulin KD. This phenomenon was more pronounced in the symbiotic strain, probably due to its generally larger cell and organelle sizes in addition to the presence of the endosymbiont (see **Fig. S4**). The red fluorescence signal of mScarlet-ETP1-expressing cells shows that although clumps of endosymbionts appear in the roundish, division-impaired host cells with reduced α-tubulin, the endosymbionts appeared to be fully divided and not filamentous, suggesting that endosymbiont division itself was not impaired **(Fig. 4F)**.

### Formation of the Z-ring at the ESDS is independent of ETP2

To assess the effects of ETP2 depletion on the localization of the bacterium-encoded FtsZ, we repeated the ETP2 KD (in cells expressing mScarlet-ETP1) followed by IFA using an antibody raised against *Ca.* K. crithidii FtsZ (Maurya et al., 2025). 24 h post-transfection, FtsZ was observed to form several foci along the filamentous endosymbionts **(Fig. 5)**, suggesting that formation of the Z-ring is independent of ETP2.

**Figure 5:**
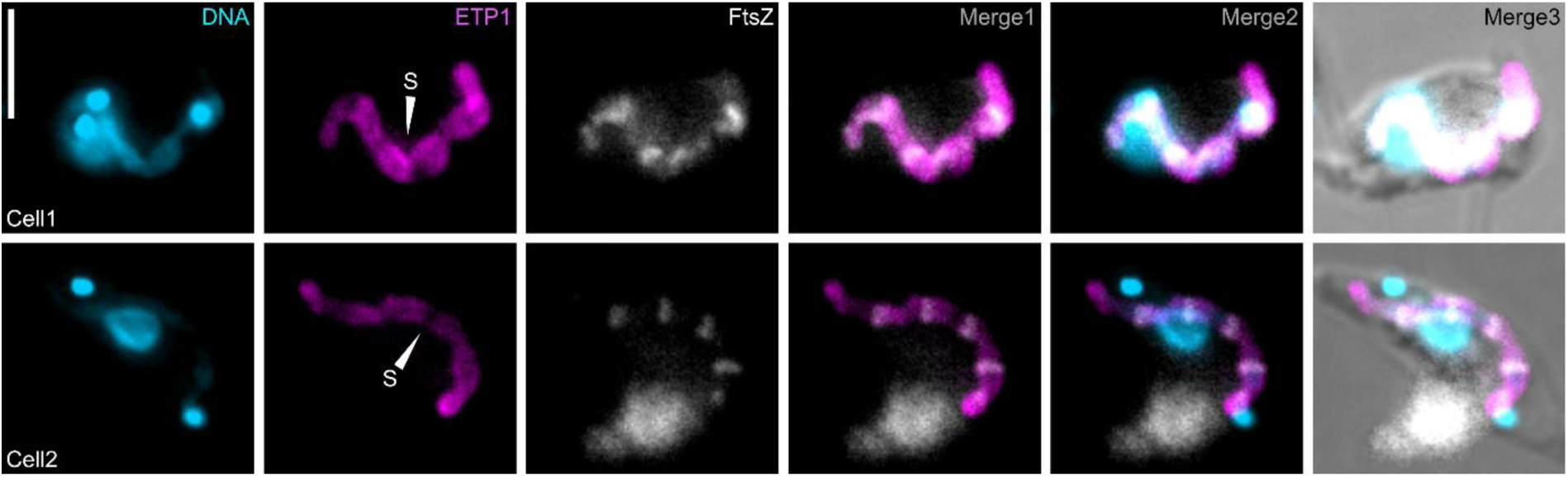
Localization of FtsZ in symbiotic *A. deanei* cells following *etp2* KD. Subcellular localization of the bacterium-encoded FtsZ in symbiotic cells expressing the endosymbiont marker mScarlet-ETP1 24 h post-transfection with MAO*_etp2_*. Shown are the fluorescence signals of Hoechst 33342-stained DNA (cyan), mScarlet-ETP1 autofluorescence (magenta), and the IFA signal for FtsZ (white). Merge1, overlay of fluorescence signals of mScarlet-ETP1 and FtsZ; Merge2, DNA, mScarlet-ETP1, and FtsZ; Merge3, DNA, mScarlet-ETP1, FtsZ, and the DIC picture. White arrowheads show long, filamentous endosymbionts (S). Scale bar is 5 µm.

### ETP2 is likely an intrinsically disordered protein and is exclusively found within the Strigomonadinae

To gain a deeper understanding of the origin and function of ETP2, we conducted BLAST searches (E-value threshold 1E-1) against both the NCBI and EukProt (Richter et al., 2022) databases. These searches yielded hits exclusively to *A. deanei* and *Strigomonas culicis* (Strigomonadinae), where the protein is annotated as ‘hypothetical protein’. To gain insights into its potential cellular function, we analyzed the ETP2 sequence for the presence of specific domains using HHPred (Söding et al., 2005) and SMART domain (Letunic et al., 2021); however, no significant hits were identified. Additionally, we employed a series of secondary structure prediction tools, including JPred4 (Drozdetskiy et al., 2015) and SPIDER3 (Heffernan et al., 2017), which predicted the presence of alpha helices and beta sheets exclusively within the C-terminal region of the sequence, beyond amino acid position 343 (**Fig. 6A**). Given the absence of predicted structural motifs in the N-terminal region, we analyzed the sequence using intrinsic disorder domain (IDD) prediction software, namely IUPred (Dosztányi et al., 2005) and SPOTD (Hanson et al., 2017). These analyses indicated a high probability of intrinsic disorder throughout the entire N-terminus, up to amino acid position 341 (**Fig. 6A**).

**Figure 6:**
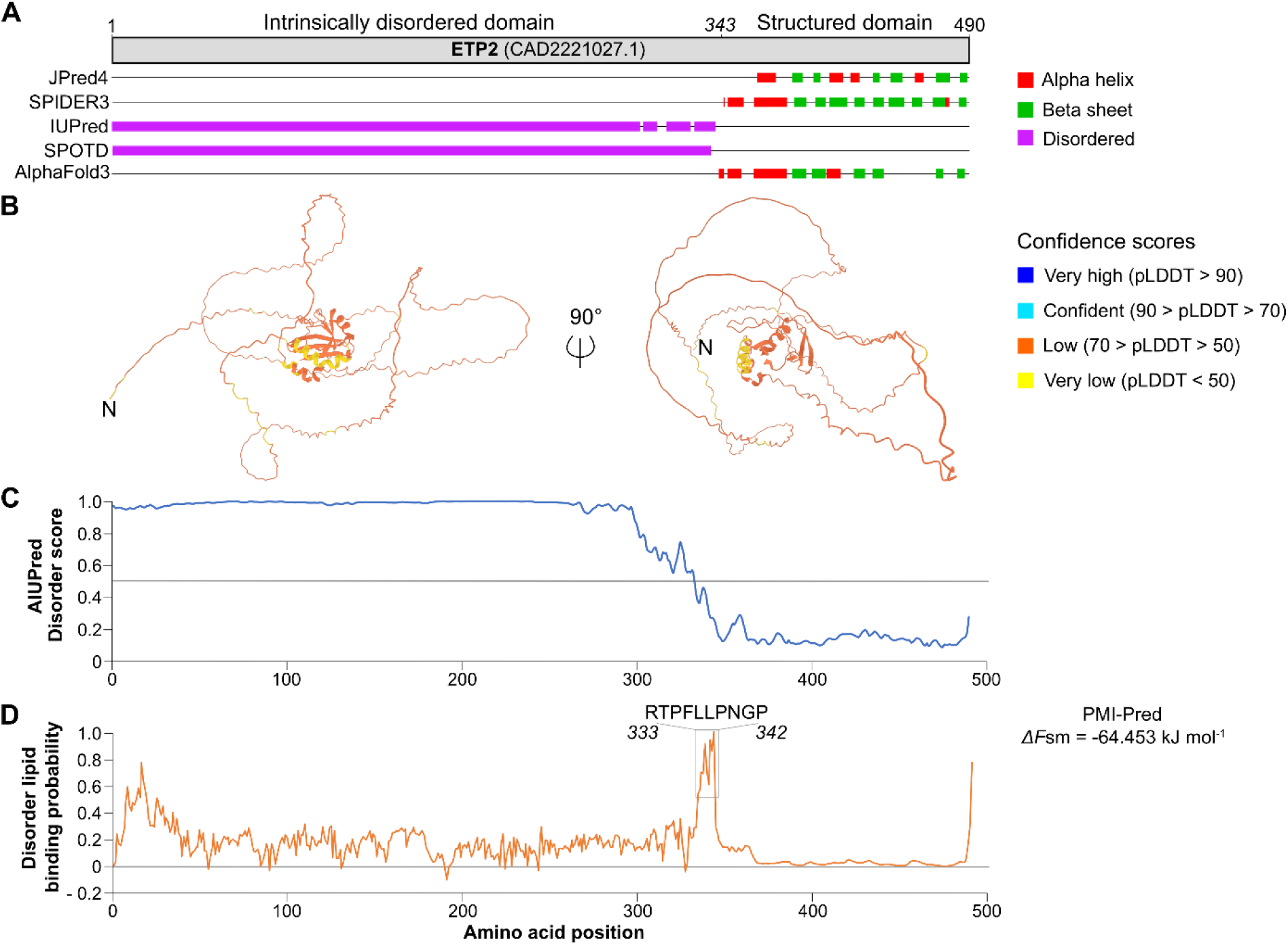
*In silico* analyses predict ETP2 as an intrinsically disordered protein with putative lipid binding capability. **A:** Overview of secondary structure elements and IDDs of ETP2 predicted by JPred4, SPIDER3, IUPred, SPOTD, and AlphaFold3 (as indicated). Consensus of the start of the structured domain of ETP2 at amino acid position 343 is indicated. **B:** AlphaFold3-predicted tertiary structure of ETP2. The N-terminus of the protein is marked with ‘N’. Confidence scores are indicated on the right. **C:** Results of AIUPred analysis of ETP2 using the default options (AIUPred – only disorder, Default smoothing). Prediction scores >0.5 (above grey line) indicate disordered regions. **D:** Results of DisoLipPred analysis of ETP2. Larger scores denote higher likelihood of a disordered lipid binding region. The region containing the highest likelihood (amino acids 333–342) is highlighted. Predicted free energy for membrane-binding of the highlighted sequence by PMI-Pred using the ‘negatively charged membrane’ option is shown on the right. Free energy values (*ΔF*sm) of <-28 kJ mol^−1^ indicate high likelihood for membrane binding.

Consistent with these sequence-based analyses, AlphaFold3 (Abramson et al., 2024) predicted for ETP2 a generally structured C-terminus and unstructured N-terminus (**Fig. 6B**). However, the predicted tertiary structure exhibited low to very low predicted local distance difference test (pLDDT) scores across the entire protein, in line with the expected confidence scores for IDDs (Ruff and Pappu, 2021). To further substantiate the prediction that the N-terminus of the protein is an IDD, we performed a more detailed analysis using AIUPred (Erdős and Dosztányi, 2024), which returned values consistent with an IDD in the N-terminal region (**Fig. 6C**).

The observed localization pattern of recombinant ETP2 suggests that the protein interacts at the ESDS with the (outer or inner) endosymbiont membrane (**Fig. 1**). Therefore, we investigated the presence of potential lipid-binding domains within the sequence by DisoLipPred (Katuwawala et al., 2022). This analysis identified a region (amino acids 333–342) with a high probability of harboring a disordered lipid-binding motif (**Fig. 6D**). The potential membrane-binding capability of this motif was further supported by PMI-Pred (van Hilten et al., 2024) analysis, which predicted a free energy of -64.453 kJ/mol for binding of this specific sequence to a negatively charged membrane, well below the required threshold of -28 kJ/mol, supporting membrane binding (van Hilten et al., 2024) (**Fig. 6D**).

## Discussion

In some associations between protists and their bacterial endosymbionts a precise coordination between host and endosymbiont cell division has evolved, ensuring a defined number of endosymbionts per host daughter cell. The molecular mechanisms underlying this cell cycle coordination are largely unknown. *A. deanei* has been recently reported to manifest nuclear control over its endosymbiont’s division by means of the DLP ETP9 that apparently forms a contractile ring structure around the ESDS and functionally compensates for the loss of essential division genes from the endosymbiont genome (Maurya et al., 2025). Here, we functionally characterized a second nucleus-encoded protein, ETP2, previously reported to localize at the ESDS in *A. deanei* (Morales et al., 2023).

Through analysis of >1000 cells expressing eGFP-tagged ETP2 (endogenously or overexpression), we found that ETP2 localizes at the ESDS specifically at cell cycle stages in which the endosymbiont divides **(Figs. 1 and 2)**. Replacement of both *etp2* alleles with the recombinant copy without a resulting phenotype **(Fig. 1C)** demonstrated that its N-terminal fusion to eGFP did not affect the localization or function of ETP2. The accumulation of ETP2 at the ESDS appears to precede the emergence of ETP9, which arrives at the ESDS after the formation of the Z-ring (compare **Fig. 2** and (Maurya et al., 2025)). The functional involvement of ETP2 in endosymbiont division is demonstrated by the striking division phenotype exhibited by homozygous *etp2* deletion mutants and by Wt cells following ETP2 KD, with filamentous endosymbionts in highly distorted host cells (**Fig. 3A-C** and **Fig. 4**). However, in both experiments numerous non-distorted host cells were also present that display Wt morphology and either harbor a single endosymbiont or lost the endosymbiont and are, hence, bound to die (**Fig. 3B-C** and **Fig. S3**). We witnessed cases where non-distorted host cells undergoing cytokinesis appeared to remain attached through their posterior ends by an undivided endosymbiont (**Fig. 3B**, white arrowhead), and apparently successfully detaching host cells not inheriting an endosymbiont (**Fig. 3B**, black arrowhead). The fact that the *etp2* deletion cell line remains alive, yet exhibits slower growth compared to the Wt (**Fig. 3D**), lends to the conclusion that despite of the disruption of the last remaining *etp2* allele, a subset of the population is still capable of undergoing cytokinesis and inheriting a morphotypical endosymbiont. Whether this outcome depends on the presence of the truncated ETP2_1-245_ that remains in the homozygous deletion mutant (**Fig. S1D**) is currently unclear.

In aposymbiotic ETP2 deletion mutants and Wt cells following ETP2 KD, no division phenotype was observed (**Figs. S2** and **4E**), demonstrating that the ETP2 function is specific for endosymbiont division and the observed distorted host morphology is a secondary effect resulting from impaired endosymbiont division. The reduced growth of aposymbiotic cells following ETP2 KD (**Fig. 4B**) may represent an off-target effect. However, in the short 5’ UTR of *etp2* no suitable binding site for a non-overlapping MAO was found that would help to verify this interpretation.

Intriguingly, in most symbiotic ETP2 deletion mutant cells, where the host is highly distorted, we observed the development of multiple kinetoplasts with an attached flagellum alongside a singular enlarged nucleus **(Fig. 3A)**. Hence, inhibition of endosymbiont division, which precedes kinetoplast division (Maurya et al., 2025), does not appear to affect kinetoplast and nucleus replication or kinetoplast segregation. In *T. brucei* and other trypanosomatids, the division furrow originates by the segregation of the flagella through separation of kinetoplasts via the basal bodies and tripartite attachment complex (TAC; a structure linking the kinetoplast and basal body), thus, making kinetoplast segregation a key initiator of cytokinesis (Kohl et al., 2003; Gluenz et al., 2011; Wheeler et al., 2013; Schneider and Ochsenreiter, 2018). Interestingly, while numerous proteins have been identified and functionally characterized that mediate this process in *T. brucei* (Schneider and Ochsenreiter, 2018), comparative genomics has indicated a potential loss of some of these regulatory genes in *A. deanei* (Cadena et al., 2024). Moreover, an additional nucleus-encoded protein that has been reported to associate with the endosymbiont, ETP5 (Morales et al., 2023), is orthologous to the KMP-11 protein, a known mediator in cytokinesis and localized to the basal bodies and flagella of *T. brucei* (Li and Wang, 2008). Whether the loss of proteins involved in kinetoplast, TAC, and basal body segregation is caused by the precedence of the endosymbiont dividing first in the *Strigomonadinae* cell cycle remains an open question.

Our BLAST, Hidden Markov Modeling, and SMART analysis results demonstrated that neither proteins similar to ETP2 nor potential domains therein are found in organisms outside of the Strigomonadinae. Secondary and tertiary structure predictions for ETP2 suggested an unstructured N-terminal domain (amino acid positions 1 – 346) ending in a potential membrane-binding motif and a structured C-terminus, however, with low confidence scores for tertiary structure prediction throughout the entire protein (**Fig. 6**). Collectively, these findings point to an origin of ETP2 as a result of *de novo* innovation within the Strigomonadinae (McLysaght and Guerzoni, 2015) and possibly its intrinsically disordered nature (Ruff and Pappu, 2021). The genesis of intrinsically disordered proteins resulting primarily from *de novo* gene innovation has been repeatedly suggested (Wilson et al., 2017; Bornberg-Bauer et al., 2021; Heames et al., 2023; Chen et al., 2024). Interestingly, due to their unstructured nature, intrinsically disordered proteins may transiently interact with multiple binding partners and, thus, act in form of signaling hubs or scaffolds as temporal organizers (Haynes et al., 2006; Patil and Nakamura, 2006; Cortese et al., 2008; Uversky, 2015; Cornish et al., 2020). Whether ETP2 is capable of acting as either a scaffold or signaling hub, recruiting other proteins involved in endosymbiont division to the ESDS, remains an exciting possibility to be explored. Since FtsZ localizes in distinct foci along the filamentous endosymbionts following *etp2* KD (**Fig. 5**), recruitment of FtsZ to the prospective ESDS, which is likely controlled by the endosymbiont-encoded Min system (Maurya et al., 2025), is apparently not affected by ETP2 depletion. Whether the recruitment of nucleus-encoded proteins such as ETP9 towards the ESDS is affected remains to be explored.

In sum, our work demonstrates that ETP2 plays a major role in the division of the endosymbiont. Additionally, we showed that disruption of endosymbiont division ultimately affects host cell division. With *A. deanei* cells containing a single, elongated endosymbiont displaying complete cytokinetic arrest alongside multiple kinetoplasts and flagella in a single host cell. Given the presence of ETP2 solely within Strigomonadinae, and lack of *in silico* identifiable homologous regions or domains, we hypothesize that ETP2 is a largely intrinsically disordered proteins resulting from *de novo* origin and may function as a scaffold or signaling hub given its early recruitment to the EDSD. Although our findings lend further credence that the endosymbiont division machinery in *A. deanei* is of dual genetic origin, the full characterization of this fascinating structure remains at its infancy.

## Methods

### Microbial strains, media, and growth conditions

Both, symbiotic *Angomonas deanei* (ATCC PRA-265) and aposymbiotic *A. deanei* (ATCC 30969) strains were grown in brain heart infusion (BHI, Sigma Aldrich) medium supplemented with 10 µg/ml hemin (Sigma Aldrich). Medium for the aposymbiotic strain was additionally supplemented with 10% v/v horse serum (Sigma Aldrich). *A. deanei* cultures were grown at 28 °C under static conditions and sub-cultured twice a week, once they reached cell densities of ∼1.0 × 10^7^ cells/ml (for the aposymbiotic strain) and ∼1.0 × 10^8^ cells/ml (for the symbiotic strain). Cell counting was performed using a Multisizer 4e cell counter (Beckman Coulter).

*Escherichia coli* TOP10 cells, used for plasmid preparation, were grown in lysogeny broth (LB) medium supplemented with 100 µg/ml ampicillin at 37 °C for 16-18 h under aerobic conditions.

### Plasmid generation

All primers used in this study are listed in **Table S1**, schematic maps of plasmids and strains used or generated in this study are displayed in **Fig. S5**.

For the generation of homozygous *etp2* deletion mutants, plasmid pAdea457 was generated. For this, the pUMA1467 backbone (Terfrüchte et al., 2014) was amplified from plasmid pAdea369 using the primer pair 3237/3244. The first half and second half of the *etp2* gene were used as 5’ and 3’ flanking region (fr) and were amplified from *A. deanei* genomic DNA (gDNA) using primer pairs 3238/3239 and 3242/3243, respectively. A fragment containing the *γ-amastin* 5’ fr with its spliced leader (SL) donor sequence, *hyg^r^*, and the intergenic region between the *gapdh1* and *gapdh2* genes, gapdh ir, was amplified from pAdea368 using primer pair 3240/3241. All the amplified fragments were assembled by Gibson cloning (Gibson et al., 2009).

For the generation of pAdea115 (used for endogenous tagging of the first *etp2* allele), the fragment *etp2* fr 3’-pUMA1467-*etp2* fr 5’ was amplified from plasmid pAdea092 using primer pair 1015/1016 and the fragment *neo^r^-*gapdh ir-*egfp*-*etp2* was amplified from pAdea035 using primer pair 1022/1023. Both fragments were ligated by Golden Gate cloning (Engler et al., 2008).

For the generation of pAdea477 (used for endogenous tagging of the second *etp2* allele), primer pair 3322/3323 was used for the amplification of the entire pAdea115 template except the *neo^r^* gene and primer pair 3320/3321 was used to amplify *hyg^r^* from pAdea260. Both fragments were ligated by Gibson cloning.

All plasmids were amplified in *E. coli* TOP10 and isolated by NucleoSpin Plasmid kit (Macherey-Nagel). Correct assembly was verified by sequencing (Microsynth AG, Balgach, Switzerland).

### Transfection of *A. deanei*

For efficient integration of a prepared DNA cassette into the *A. deanei* nuclear genome, ∼40 µg of plasmid DNA was digested with restriction enzyme(s) cutting at the 5’ and 3’ ends of the insertion cassette (see maps in **Fig. S5**). The linearized cassette was transfected into *A. deanei* cells as described earlier (Morales et al., 2023). In brief, 1.0 × 10^7^ cells were harvested and resuspended in 18 µl of P3 primary cell solution. 2 µl of the digested plasmid (2-4 µg DNA) were mixed with the cells and pulsed with the program FP158 using Nucleofector 4D (Lonza). Cells were immediately transferred to 5 ml growth medium without antibiotics. Cells were allowed to recover for 6 h at 28 °C. After this incubation, cells were diluted 10-fold in the same medium with the respective selection antibiotic(s) added at final concentrations of 500 µg/ml for both G418 (Sigma Aldrich) and hygromycin B gold (InvivoGen). Diluted cells were distributed in 200 µl aliquots on a 96-well plate and incubated at 28 ^°^C until single wells showed turbidity indicating the growth of clonal cell lines (typically 5-6 days). Clones were isolated and used further for verifications by PCR.

### Verification of transgenic cell lines by PCR

To verify selected clones by PCR, gDNA was isolated from cells using 200 µl of DNAzol (Thermo Fisher) according to the manufacturer’s instructions and dissolved in 20 µl of sterile dH_2_O. Primers that were used **(see Table S1)**, either binding in the genomic region outside of the inserted cassette or one primer binding outside and the other inside the cassette. Only verified strains were used for further analysis.

### Southern blot analysis

For the replacement of the second *etp2* allele (that caused a phenotype in the context of *etp2* disruption, but not for *egfp* tagging), insertion of a single cassette was verified by Southern blot analysis as described before (Morales et al., 2023). Briefly, ∼7.5 µg of gDNA was isolated by the DNeasy Blood & Tissue kit (Qiagen), digested overnight using appropriate restriction enzymes, and resolved on 0.8% w/v agarose gel. Next, DNA was transferred to a nylon membrane (Nytran N Nylon Blotting Membrane, 0.45 μm; GE Healthcare Life Sciences) overnight by capillary method and fixed on the membrane using 0.120 Joules with a UV-strato-Crosslinker (Analytik Jena). A digoxigenin (DIG)-labelled probe against *hyg^r^* (*α-hyg^r^*; see **Fig. S1**) was generated using the primers indicated in **Table S1** and hybridized to the DNA immobilized on the membrane at a hybridization temperature of 53 °C. Lastly, the membrane was developed as per the protocol supplied with the DIG-High Prime DNA Labeling and Detection Starter kit II (Roche Applied Science). The resulting chemiluminescence signal was detected by a Chemidoc MP (Bio-Rad).

### Epifluorescence microscopy

For visualizing cells, epifluorescence microscopy was performed as described before (Morales et al., 2023). In brief, cells in the mid-log phase were mixed with formaldehyde (FA) to a final concentration of 4% w/v and incubated in dark at room temperature (RT) for 10-15 min. Next, cells were washed twice and resuspended in phosphate buffered saline (PBS). Then, cells were spotted onto a glass slide coated with 0.01% poly-L-lysine (Specialty Media), mixed with 30 µg/ml Hoechst 33342 in PBS, and incubated at RT in the dark for 10-15 min. After incubation, cells were washed three times with PBS and coated with antifade reagent SlowFade Diamond (Thermo Fisher Scientific). Images were acquired with an Axio Imager M.1 (Zeiss, Oberkochen, Germany) using an EC Plan-Neofluar 40x/1.3 oil Ph3 or 100x/1.30 Oil Ph3M27 objective (Zeiss). Images were analyzed with Zen Blue v2.5 and processed with ImageJ v2.0 software.

### Immunofluorescence assay (IFA)

For visualizing FtsZ, IFA was performed as described before (Maurya et al., 2025). All steps were performed at RT in the dark. In brief, cells in the mid-log phase were mixed with FA to a final concentration of 4% w/v and incubated for 15 min. Cells were collected, washed with PBS, and resuspended in an appropriate volume of PBS. Then, 20 µl cell suspension was spotted onto 0.01% poly-L-lysine-coated glass slides and incubated for 20 min. After incubation, unbound cells were removed and attached cells permeabilized with 10 µl 0.2% v/v Triton-X100 in PBS for 20 min. Then, cells were washed in PBS and incubated with 1% w/v blocking solution (BS; albumin bovine fraction V, pH-7.0, Serva) in PBS for 45 min. Next, 15 µl of 1:80 diluted anti-FtsZ_pepC_ primary antibody (raised in rabbit against *Ca.* K. crithidii FtsZ, see (Maurya et al., 2025)) in 1% BS was applied and incubated for 1.5 h followed by three washes in 1% BS. The 1:500 diluted anti-rabbit IgG (goat, polyclonal, Abberior Star-red, Ex: 638 nm) secondary antibody in 1% BS was applied and incubated for 1 h followed by three washes in 1% BS. After antibody staining, DNA was stained with Hoechst 33342. Finally, cells were coated with 7 µl antifade reagent SlowFade Diamond (Thermo Fisher Scientific) and imaged with an Axio Imager M.1 as described before. mScarlet-ETP1 and eGFP-ETP2 were detected based on their autofluorescence. For reconstruction of cell cycle stages, more than 1,000 cells were analyzed with Zen Blue v2.5 software.

### Fluorescence *in situ* hybridization (FISH)

For visualizing the endosymbionts by FISH, cells from mid-log phase were mixed with FA to a final concentration of 4% w/v and incubated for 30 min. Cells were harvested thereafter, washed 3 times with PBS, and resuspended in an appropriate volume of PBS. Then, 20 µl cell suspension was spotted onto 0.01% poly-L-lysine-coated glass slides and air-dried for 30 min. The slides were consecutively placed in 50%, 80%, and 100% ethanol for 3 min each to dehydrate the cells. Then, 5 µl of a 5’ Cy3-labeled Eub338 probe (50 ng DNA µl^−1^) directed against the bacterial 16S rRNA (Amann et al., 1991; Fuchs et al., 2007) was mixed with 45 µl of hybridization buffer (900 mM NaCl, 20 mM Tris-HCl pH 7.2, 35% formamide, 10% SDS in dH_2_O) and spotted onto the dry slides. The slides were incubated at 46 °C for 2 h in a humid chamber. Afterwards, the slides were washed in FISH-wash buffer (80 mM NaCl, 20 mM Tris-HCl pH 7.2, 10% SDS in dH_2_O) and incubated at 48 °C for 25 min. The slides were rinsed twice with dH_2_O and mixed with 30 µg/ml Hoechst 33342 in PBS, and incubated at RT in the dark for 10-15 min. After incubation, cells were washed three times with PBS and coated with SlowFade Diamond (Thermo Fisher Scientific). Images were acquired and analyzed as described before.

### Confocal microscopy

Slides were prepared as described earlier (Maurya et al., 2025). The protocol was the same as for epifluorescence microscopy slide preparation except that a high precision coverslip (LxW-22×22 mm, No. 1.5H, thickness-170 µm, Marienfeld, VWR) was used to spot 0.01% poly-L-lysine. All steps were performed on this coverslip. Later, the coverslip was put upside down on a glass slide. Imaging was performed at an inverted confocal microscope, Leica TCS SP8 (STED 3X), with standard settings as described before (Morales et al., 2023) except that the laser line was set at 35% for eGFP and 15% for mScarlet. For deconvolution of the raw data, Huygens Professional v.23.10.0p7 64b was used with default settings except that in manual mode a threshold for background extraction was set based on background fluorescence signals. Generation of supplementary movies was performed as described earlier (Maurya et al., 2025).

### KD assays using MAOs

For KDs, *A. deanei* cells were transfected with MAOs as described earlier (Maurya et al., 2025) using MAO*_etp2_* (5’-CGTAGTCCATTTTGGTGTGTATGAT-3’) or MAO*_tub_* (5’-TGAATGCAGATAGCCTCACGCATGG -3’, as a positive control), all synthesized by Gene Tools, LLC, Philomath, OR, USA. After transfection, cells were transferred into BHI medium supplemented with hemin and horse serum without antibiotics for all strains and incubated at 28 °C. Samples were collected at 6 h, 12 h, and 24 h post-transfection and used for Hoechst 33342 staining and epifluorescence microscopy as described before.

### Growth measurements

For the homozygous *etp2* mutant cell line, growth analysis was performed as described before (Maurya et al., 2025). Briefly, 4 × 10^5^ cells/ml were inoculated in triplicates in 5 ml BHI medium supplemented with hemin without antibiotics and incubated at 28 °C. Cells were counted in triplicates at 24 h and 48 h using a cell counter (Multisizer 4e, Beckman Coulter). For MAO-treated cells, counting was performed in triplicates at 6 h, 12 h, and 24 h post-transfection.

### *A. deanei* cell and organelle measurements

For measuring the width (W) and length (L) of symbiotic and aposymbiotic *A. deanei* strains, cell line Adea126 expressing mScarlet-ETP1 and Wt aposymbiotic strains were used, respectively. For W and L measurements of the cell, DIC images were used. For kinetoplast (K) and nucleus (N), Hoechst 33342 staining was used. For measuring the endosymbiont (ES), mScarlet-ETP1, localized at the bacterial envelope, was used. Measurements were performed manually with Zen v.2.5 on cells that were categorized into different cell cycle stages. Mean values were calculated and plotted using GraphPad Prism v5.0.

## Acknowledgements

This study was supported by the PhD fellowships of the Jürgen Manchot Graduate School ‘‘Molecules of Infection’’, MOI III (to GE), MOI IV (to AKM), and MOI V (to LRC). Additionally, we thank Dr. Sebastian Hänsch from the Center for Advanced Imaging (Cai - Heinrich Heine University Düsseldorf) for help with confocal microscopy, leading to movies S1 and S2.

## Author contributions

ECMN and AKM designed the research. AKM, LRC, and GE performed the experiments. All authors analyzed the data. ECMN supervised the research. AKM, LRC, and ECMN wrote the manuscript.

## Supplementary Materials

**Figure S1:**
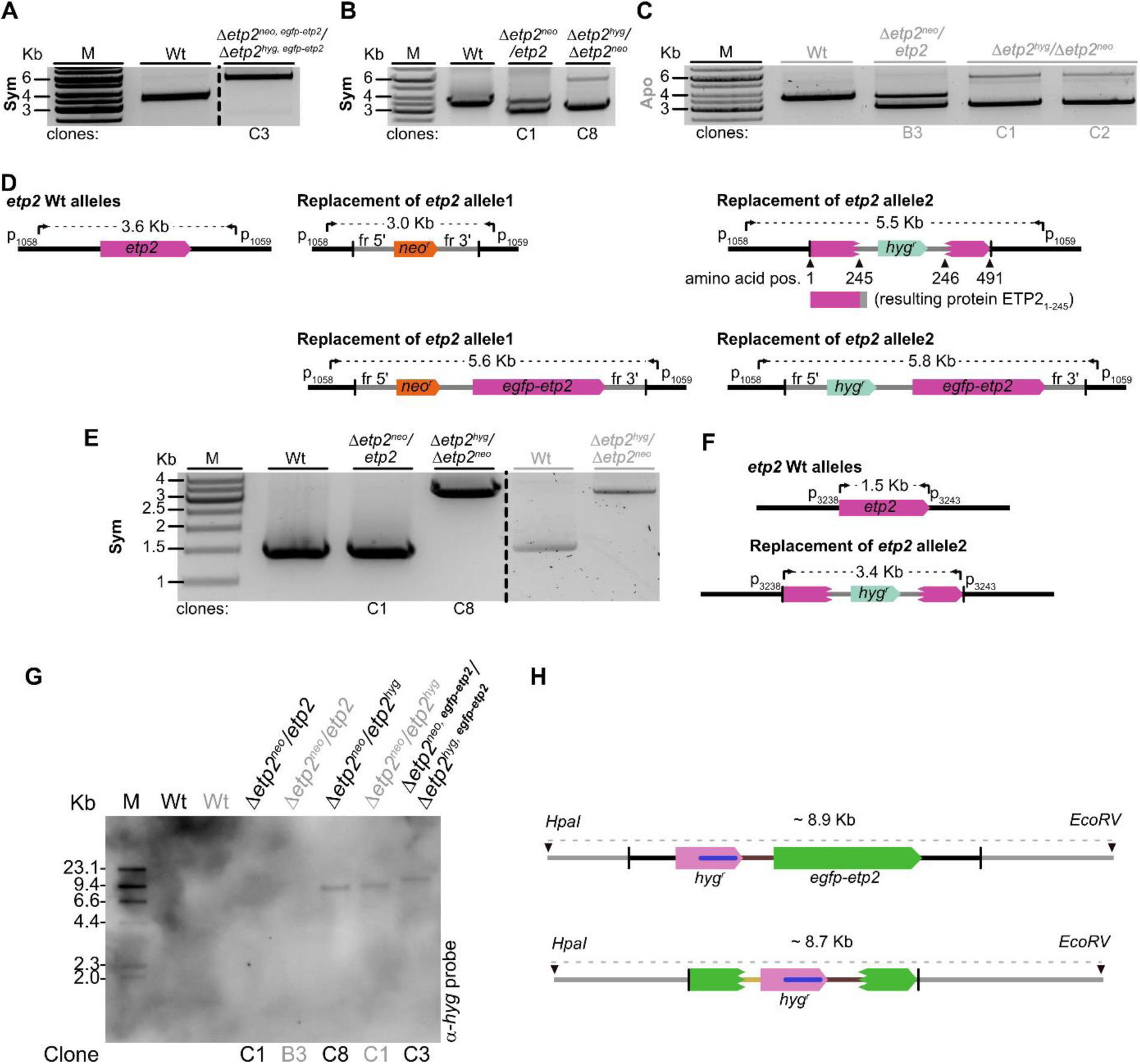
Verification of recombinant *A. deanei* cell lines by PCR and Southern blot analysis. **A:** Verification of the symbiotic strain with both alleles of *etp2* replaced by an *egfp-etp2*-containing cassette. **B:** Verification of symbiotic *etp2* heterozygous and homozygous deletion mutant cell lines. **C:** Verification of aposymbiotic heterozygous and homozygous *etp2* deletion mutant cell lines. **D:** Maps of Wt and modified *etp2* loci with expected band sizes detected by PCR (for A-C). Black arrowhead marks the amino acid position. **E:** PCR analysis of presence/absence of an intact *etp2* copy in symbiotic and aposymbiotic *A. deanei* Wt and *etp2* deletion mutant cell lines. **F:** Maps of Wt and modified *etp2* locus with expected band sizes detected by PCR in E. Black text shows symbiotic and grey aposymbiotic strains. Dotted lines show expected PCR band sizes. Curved arrows show primer binding sites. Vertical black lines on the maps show borders of inserted DNA fragments in the genome. Broken pink/green boxes on the maps show first and second halves of the *etp2* ORF used as fr 5’ and fr 3’ for homologous recombination, respectively (in D, F, H). **G:** Southern blot verification of *A. deanei* strains with Wt and modified *etp2* loci using a probe against *hyg^r^*. M, DIG-labelled DNA marker. **H:** Maps of Wt and recombinant *etp2* loci analyzed by Southern blot. Black arrowheads on the maps indicate the cutting site for the indicated restriction enzymes. The grey dotted lines indicate the expected size of the digested fragment. Binding sites of the α-*hyg* probe are marked by a blue bar. Abbreviations: fr, flanking region; *hyg^r^*, hygromycin resistance gene, and *neo^r^*, neomycin resistance gene.

**Figure S2:**
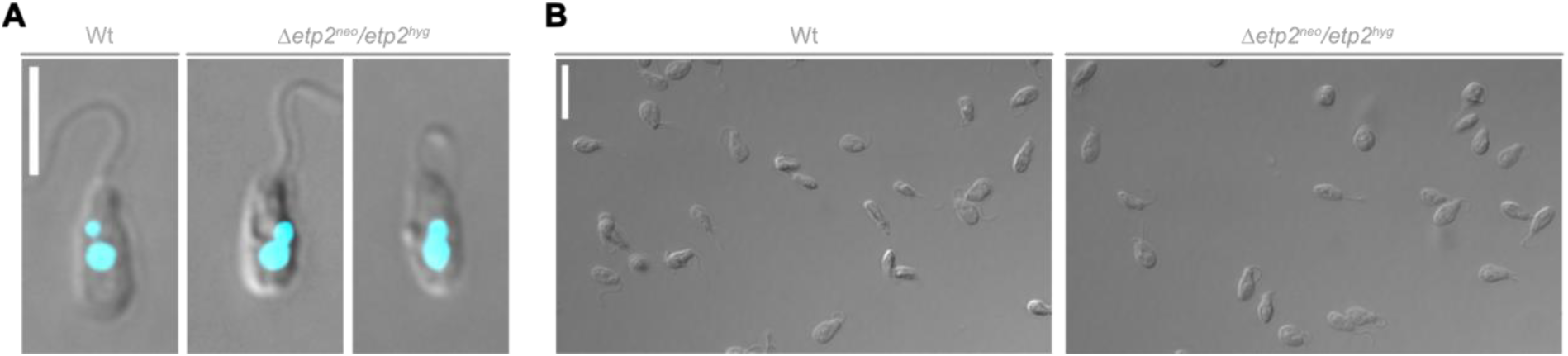
Effects of deletion of both alleles of *etp2* in the aposymbiotic *A. deanei* strain. **A:** The figure shows representative images of Wt and the homozygous *etp2* deletion mutant cell line in the aposymbiotic *A. deanei* strain. Images show an overlay of DIC and Hoechst 33342 fluorescence (cyan). Scale bar is 5 µm. **B:** Overview DIC images of cells of the same cultures. Scale bar: 10 µm. Grey text/bars indicate aposymbiotic strains.

**Figure S3:**
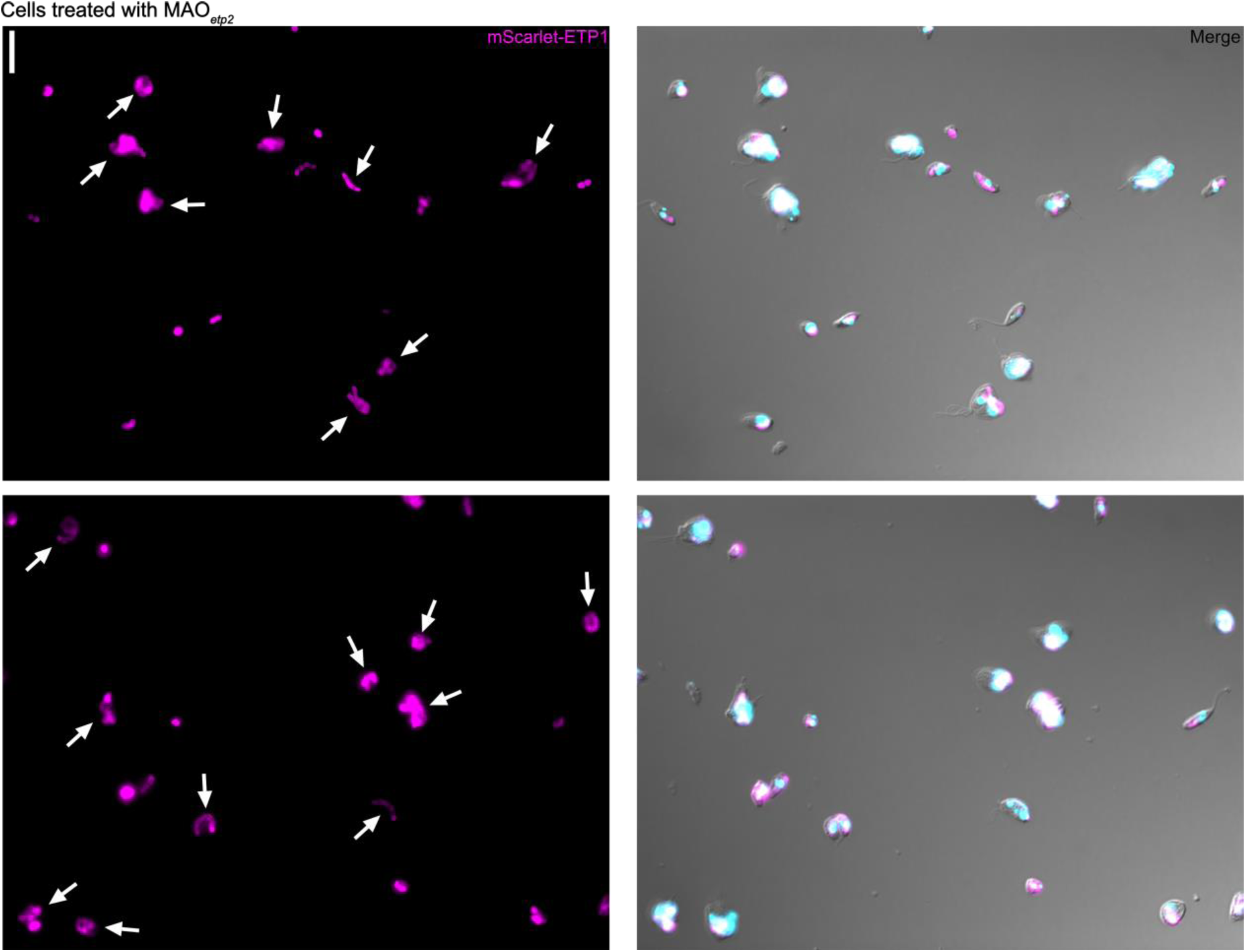
Effects of KD of ETP2. *A. deanei* cells expressing the endosymbiont marker mScarlet-ETP1 24 h post-transfection with MAO*_etp2_*. Shown are the mScarlet channel alone (on the left) and a merge of the mScarlet and Hoechst 33342 channels with the DIC image (on the right). Arrows show filamentous endosymbionts. Scale bars are 10 µm.

**Figure S4:**
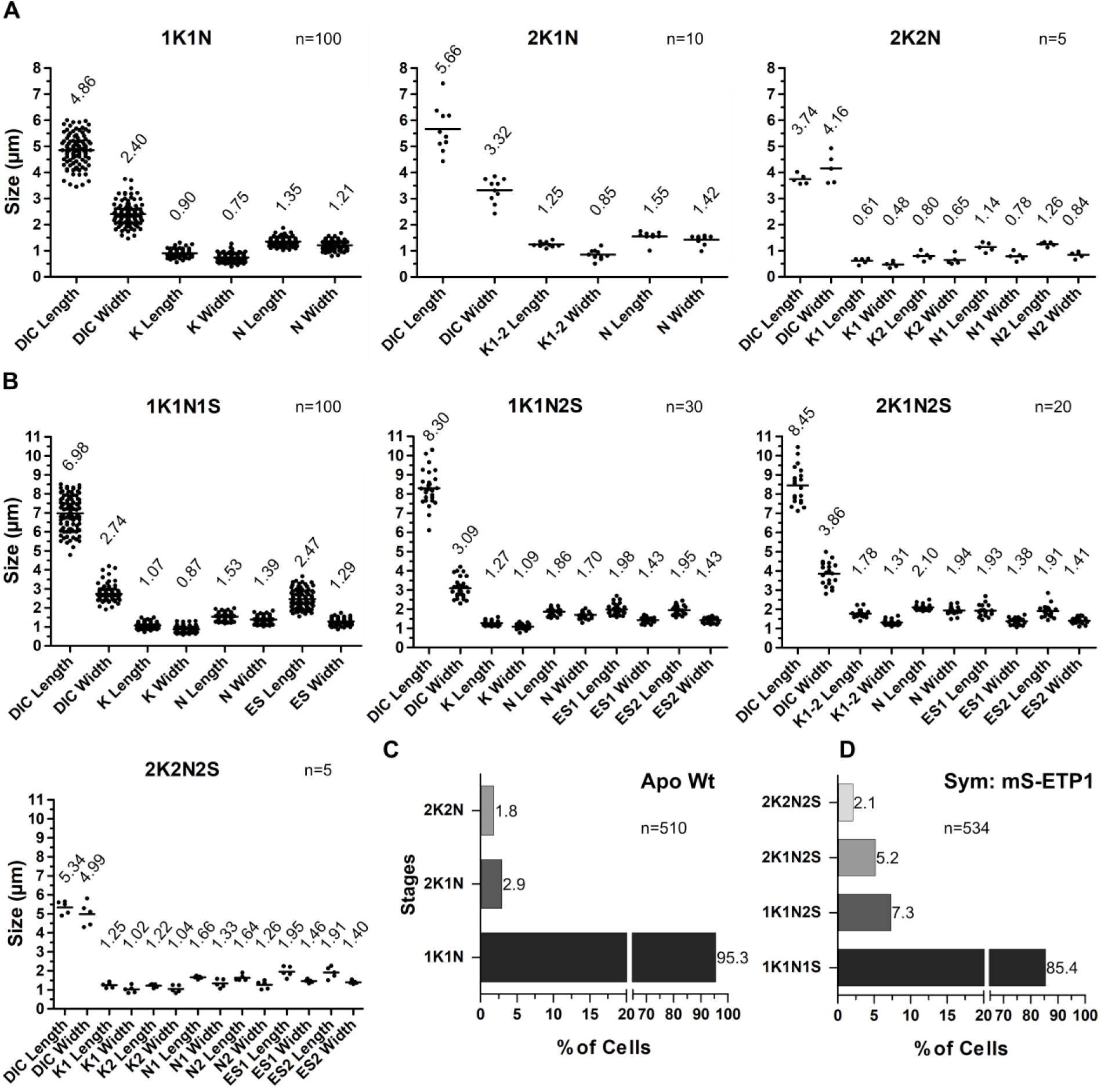
Measurements of cell and organelle sizes for symbiotic and aposymbiotic strains. **A:** Measurements for aposymbiotic Wt cells at different cell cycle stages-1K1N, 2K1N, and 2K2N (see below). **B:** Measurements for symbiotic cells expressing the endosymbiont marker mScarlet-ETP1 at different cell cycle stages-1K1N1S, 1K1N2S, 2K1N2S, and 2K2N2S (see below). Mean values are represented on each plot for each measurement by a horizontal line. DIC images were used to measure cell size, Hoechst 33342-stained DNA for kinetoplast (K) and nucleus (N), and mScarlet-ETP1-labelled endosymbiont (ES) for bacterium measurements, respectively. K1-2 represents two kinetoplasts that are still not completely separated. **C:** Quantification of approximately 500 aposymbiotic Wt cells categorized into different cell cycle stages. **D:** Quantification of approximately 500 symbiotic cells expressing mScarlet-ETP1 categorized into different cell cycle stages. Cell cycle stages are described by the number of kinetoplasts (K), nuclei (N), and endosymbionts (S) per cell.

**Figure S5:**
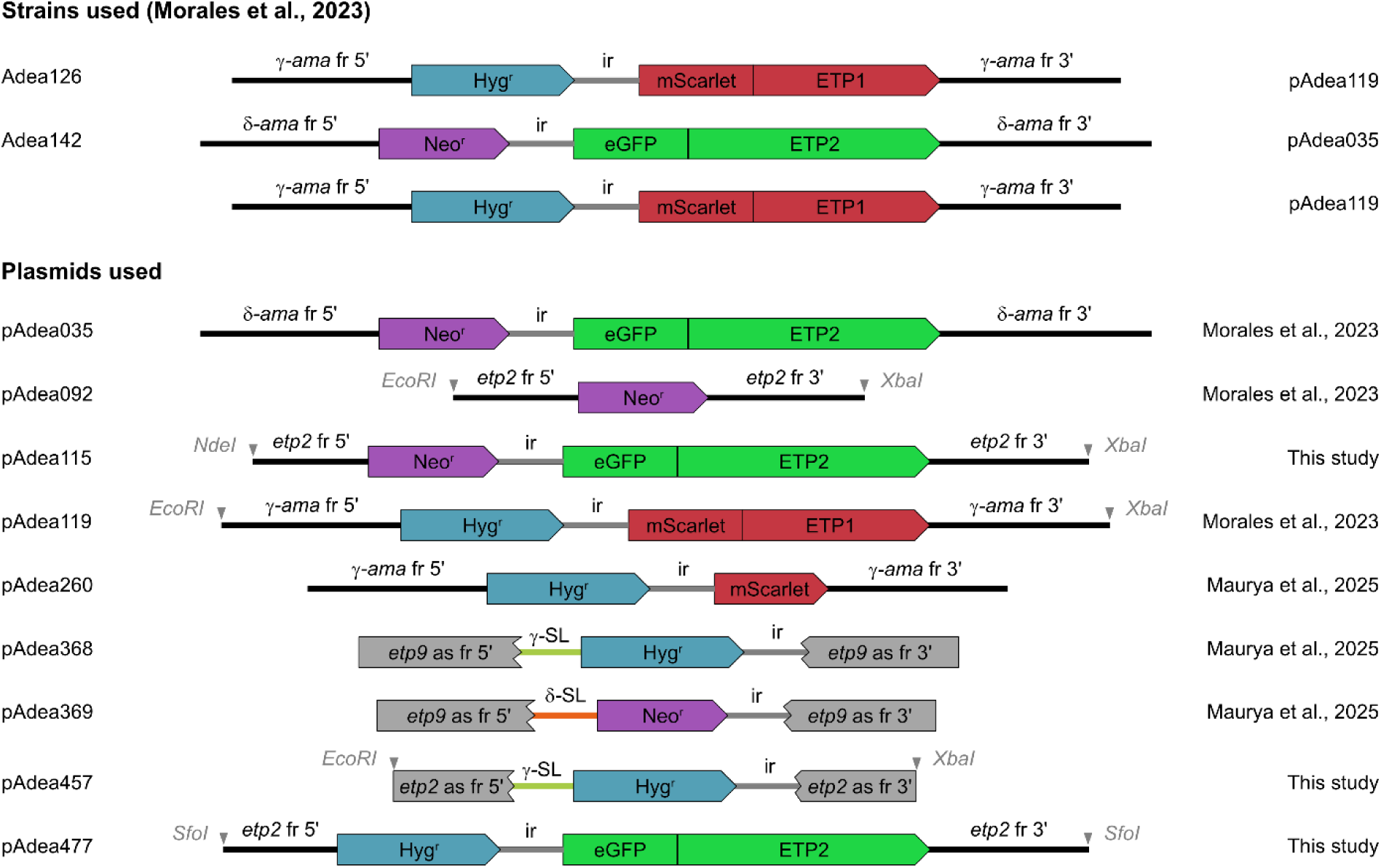
Schematic maps of all plasmids and *A. deanei* strains used in this study. Previously generated plasmids such as pAdea092 and pAdea119 were used in this study to generate new Adea strains or as templates to generate new plasmids in this study (pAdea260, pAdea368, and pAdea369). All other plasmids were generated in this study. Abbreviations: fr, flanking region (∼0.7 - ∼1.0 Kbp used for homologous recombination); ir, intergenic region between *gapdh I* and *gapdh II* genes; δ/γ*-ama*, δ- and γ-*amastin* gene (for nomenclature see Morales et al. 2016); *etp2* fr 5’, upstream flanking region of *etp2* gene; *etp2* fr 3’, downstream flanking region of *etp2* gene; *etp2/9* as fr 5’, first half of the respective *etp* ORF used as 5’ flaking region; *etp2/9* as fr 3’, second half of the ORF used as 3’ flaking region; δ/γ-SL, 5’ flanking region of the δ/γ-*amastin* genes, respectively including the spliced leader acceptor sequence; Neo^r^/Hyg^r^, neomycin and hygromycin resistance markers encoded by *neo^r^*/*hyg^r^* genes, respectively. Arrowheads show approximate cutting sites of restriction enzymes on the plasmid used for linearization before transfection.

**Table S1:**
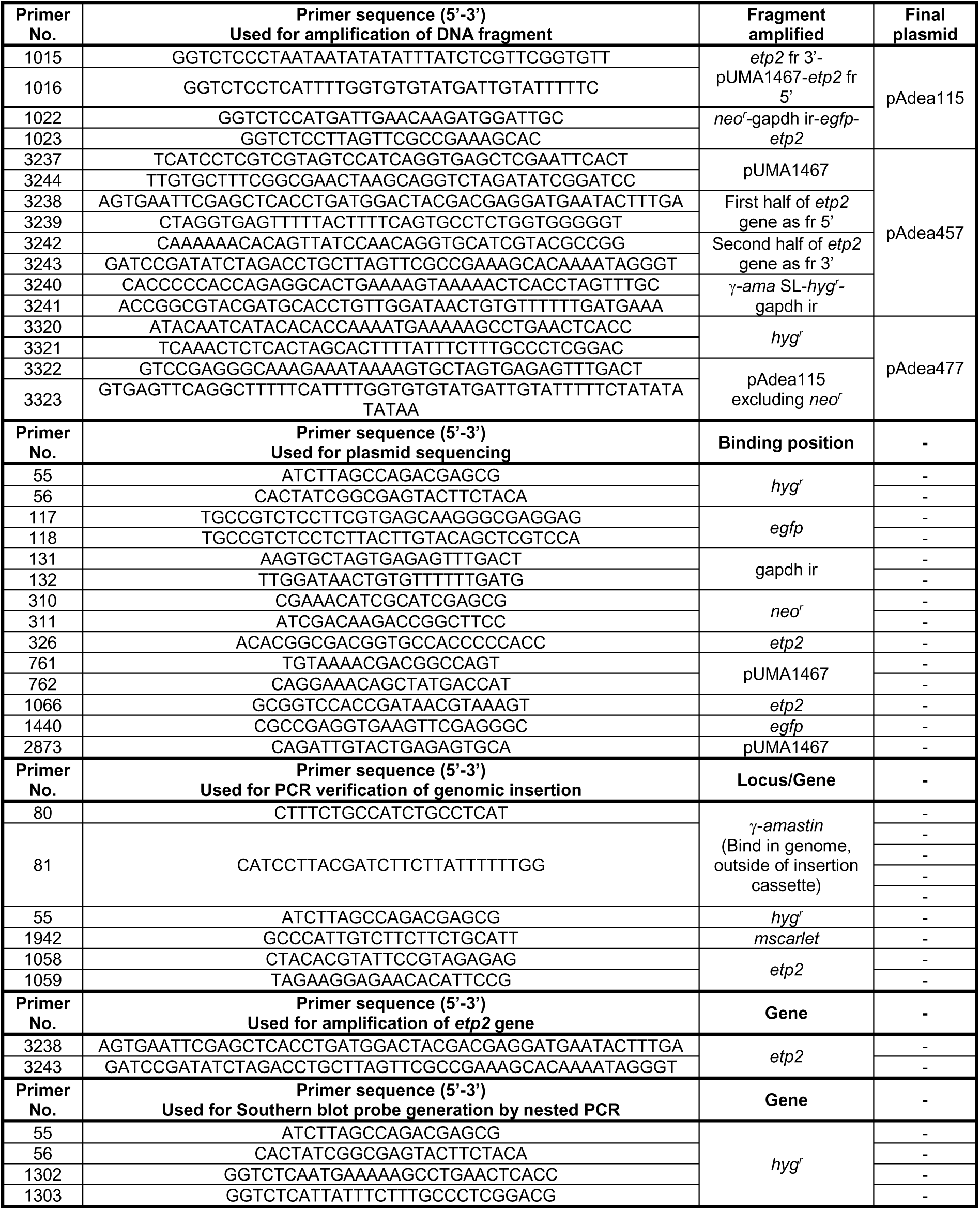
All primers used in this study. Primers for the generation of plasmids, plasmid sequencing, verification of genomic insertions, and generation of Southern blot probes are described below.

## Supplementary movies

**Movie S1:**

Cell co-expressing eGFP-ETP2 (Δδ*-ama^egfp-etp2^*) and mScarlet-ETP1 (Δγ*-ama^mS-etp1^*) that is at an early stage of endosymbiont division showing a ring-shaped fluorescence signal around the ESDS when scrolling through confocal Z-stacks. Shown are individual channels for Hoechst 33342, mScarlet, and eGFP fluorescence, as well as a merge of all channels. Scale bar: 2 µm.

**Movie S2:**

Cell co-expressing eGFP-ETP2 and mScarlet-ETP1 that is at a later stage of endosymbiont division showing an x-like fluorescence signal at the ESDS when scrolling through confocal Z-stacks. Shown are individual channels for Hoechst 33342, mScarlet, and eGFP fluorescence, as well as a merge of all channels. Scale bar: 2 µm.

